# Refinement of the *Brassica napus* NLRome using RenSeq

**DOI:** 10.1101/2024.07.24.604657

**Authors:** Jiaxu Wu, Soham Mukhopadhyay, Edel Pérez-López

**Author notes:** These authors contributed equally to this work (Jiaxu Wu, Soham Mukhopadhyay).

## Abstract

Canola (*Brassica napus* L.), a valuable oilseed crop, faces significant challenges from diseases such as clubroot and blackleg, threatening global yields. To improve disease resistance, it’s crucial to accurately annotate nucleotide-binding leucine-rich repeat receptors (NLRs), which are key components of plant immune systems. This study focuses on refining the NLR repertoire (NLRome) of the Westar cultivar using Resistance gene enrichment sequencing (RenSeq). Initially, only 345 NLR genes were identified in the Westar genome annotation, a figure significantly lower than other canola cultivars. By employing RenSeq, we expanded the annotated NLR genes to 715, including 287 full NLRs and 428 partial NLRs, thus providing a more comprehensive understanding of the NLR diversity in this cultivar. Our findings reveal crucial genomic regions with previously misannotated NLR genes and identify key integrated domains that may contribute to enhanced disease resistance. This refined NLRome not only advances the fundamental understanding of *B. napus* genetics but also offers valuable insights for breeding programs and biotechnological applications aimed at improving crop resilience. The data generated in this study is available in public repositories for further research and application.

---

Canola (*Brassica napus* L.) is primarily cultivated as an oilseed crop with significant economic value (Javed et al., 2023). However, the prevalence of diseases such as clubroot, sclerotinia stem rot, and blackleg threatens the canola industry worldwide (Chen et al., 2021). Research and breeding efforts often focus on identifying novel resistance genes (R genes) involving receptor-like proteins and kinases (RLPs and RLKs), and nucleotide-binding and leucine-rich repeat (NLR) receptors to understand the resistance mechanisms and control the spreading of plant diseases. NLR receptors, which are highly expanded and diverse in the plant genome, play a crucial role in initiating a robust immune response upon perceiving pathogen-secreted effectors (Kourelis et al., 2021; Thomas et al., 2024). Canonical NLRs in plants consist of three domains including an N-terminal domain, a central nucleotide-binding (NB-ARC) domain, and a C-terminal leucine-rich repeat (LRR) domain. An NLR is usually classified based on its N-terminal domain as Toll/interleukin 1 receptor (TIR) NLR (TNL), coiled-coil NLR (CNL), and Resistance to powdery Mildew 8 (RPW8) coiled-coil NLR (RNL) (Marchal et al., 2022). It is reported that the activation of these N-terminal domains of NLR can lead to programmed cell death in plant tissues known as the hypersensitive response (HR) to restrict the pathogen infection (Kourelis et al., 2021). Accurate annotation of NLR gene repertoire is essential for further understanding their functions and engineering resistant crops (Jupe et al., 2013).

The *B. napus* Westar cultivar (hereafter Westar canola) is a spring-type, low erucic acid cultivar that is widely used in genetic mapping and transformation studies. It also serves as a universal susceptible host standard for studying canola disease resistance to pathogens (Javed et al., 2023). A long-read-based high-quality genome was released for Westar canola, which is predicted to carry 97,514 annotated genes (https://yanglab.hzau.edu.cn/BnIR/germplasm_info?id=Westar.v0). While studying the NLR repertory of Westar canola based on the available gene annotation file, we found the presence of only 345 genes with domain signatures associated with NLR proteins using InterproScan 5 and NLRtracker (an NLR annotation tool) (Supplementary Table S1) (Kourelis et al., 2021). This number is very low compared to other *B. napus* canola cultivars such as ‘ZS11’ (Genome Warehouse Accession number: GWHANRE00000000) and ‘Darmor’ (NCBI Accession number: GCF_020379485.1) which are predicted to carry a total of 597 and 641 NLR loci, respectively (Alamery et al., 2018; Chen et al., 2021). This made us wonder if (*i*) the Westar canola genome was properly annotated, and if (*ii*) we could further refine the canola NLRome. To answer those questions, we produced the first canola NLRome using the Westar canola genome and resistance gene enrichment sequencing (RenSeq), a method that has advanced R genes discovery and cloning in crops like wheat and potato (Jupe et al., 2013). RenSeq is a targeted sequencing method that enriches and sequences NLRs within a plant genome, offering a less expensive and less laborious alternative for NLR genetic mapping or NLR discovery studies (Jupe et al., 2013).

In this study, a set of 74,738 RenSeq baits was designed based on the ‘ZS11’ canola genome NLR nucleotide sequences (Genome Warehouse accession number: GWHANRE00000000) by Arbor Bioscience (Fig. 1a, Figure S1, Supplementary File 1) (Chen et al., 2021). We used 80 nt probes and 3x tilling with a total target size of 2,951,019nt and an average GC content of 37.6%. Targets were softmasked for repeats against the eudicot database. Strings of Ns 1-10 nt long were replaced with T. High molecular weight genomic DNA was extracted from 3-week-old homozygous doubled-haploid Westar canola seedlings with a CTAB method as previously described (Jupe et al., 2013). Illumina library enrichment and NLR capture sequencing were performed by myBaits^®^ Custom Hybridization Capture Kits (Daicel Arbor Biosciences, MI, USA) following their standard protocols, resulting in 45 million 150 bp paired-end Illumina reads (Bioproject: PRJNA1137270). Next, we mapped the reads to the Westar genome using BWA-MEM2 and visualized them using Geneious Prime (https://www.geneious.com/). In parallel, the NLR-Annotator (Steuernagel et al., 2020) was used to predict NLR loci across the genome.

**Figure 1.**
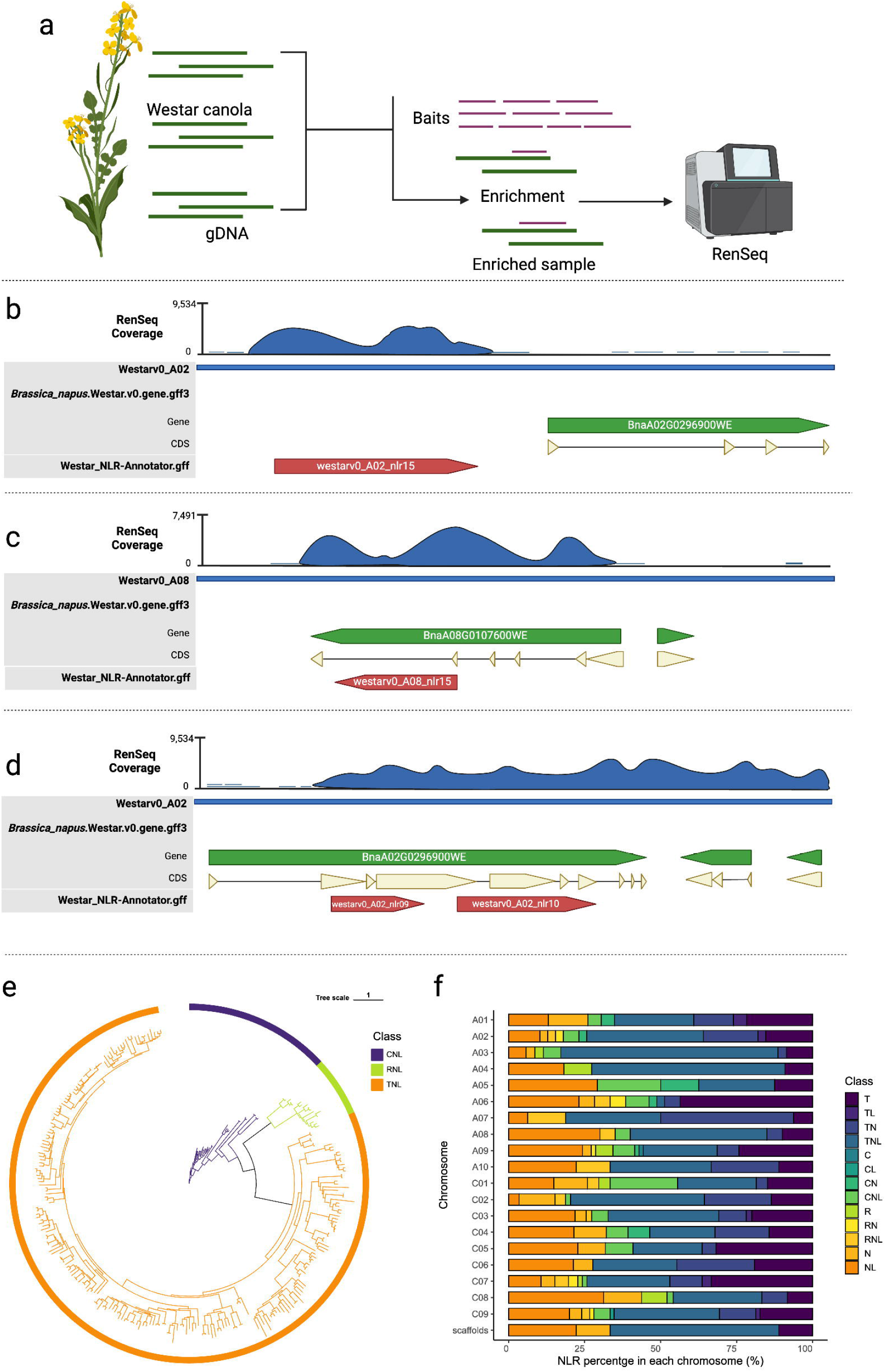
*Brassica napus* NLRome refinement using RenSeq. **a**, schematic representation of the workflow followed in this study. Full representation can be found in Supplementary Fig. S1. **b**, example of NLR missed in annotation file. **c**, example of NLR wrongly annotated in the annotation file. Here we present the case of *Crr1a*, a well-known clubroot resistant gene that, although present in the Westar canola genome, is not functional. **d**, example of multiple NLRs annotated as a unique NLR in the reference annotation file. **e**, phylogenetic representation of the 287 full NLRs identified in the Westar genome. Clades with different colors represent different types of NLR genes (TNL, CNL and RNL). These sequences were aligned using Muscle V5. Maximum likelihood analysis was conducted by IQ-TREE2 software, bootstrapped 1000 times, tree scale = 1. **f**, Frequence of NLR classes in genomes A and C by chromosomes.

Using Geneious software, we overlaid the NLR-Annotator prediction and the gene annotation supplied with the genome (Supplementary File S2) on top of the BWA-MEM2 mapping. This allowed us to manually inspect the genes that are correctly annotated with sufficient RenSeq read depth (at least 50x) and NLR-Annotator coverage. All 345 genes that were previously identified by InterproScan and NLRtracker and annotated in the Westar canola genome were also identified using this methodology (Supplementary Fig S2, Table S1 and File S2).

Wrongly annotated NLRs in the Westar genome fell into three main categories: 1) predicted NLR-encoding genes absent in the annotation file; 2) wrongly annotated NLR-encoding genes in the annotation file; and 3) annotated NLR-encoding genes in the annotation file that contained two or more NLRs (Fig. 1b-d). For example, we found the gene model of the *Crr1a* allele, a known clubroot R gene originally found in resistant *B. rapa* cultivar ‘G004’, to be wrongly annotated in the Westar genome (Fig 1c). To correctly annotate the remaining positions where RenSeq read and NLR-Annotator coverages were present, but gene annotations were either missing or partial, we wrote a custom script to extract such regions with 1 Kb flanking positions and with at least 50x ResSeq read depth (https://github.com/Edelab). AUGUSTUS (https://bioinf.uni-greifswald.de/augustus/) was used to *de-novo* predict genes in the extracted regions and the longest isoform was selected for further analysis. Using InterproScan with the Pfam database we found that 370 of those sequences were carrying NLR-associated domains. Thus, by combining RenSeq mapping, the NLR-Annotator annotation, and de-novo gene prediction of problematic regions using AUGUSTUS, we increased the number of correctly annotated NLR genes from 345 to 715 (Supplementary File S3). NLRtracker was used to classify the 715 amino acid sequences based on their domain composition, identifying 287 full NLRs and 428 partial NLRs in the Westar genome, diversity that was confirmed through a maximum likelihood phylogenetic analysis using IQ-TREE2 (Fig. 1e, Supplementary Table S1). Chromosomes C09 and A09 were found to have the most abundant NLRs, with 75 and 68, respectively (Fig. 1f, Supplementary Table S2). However, only 9 NLRs are present on chromosome A10 (Fig. 1f, Supplementary Table S2). Among the full NLRs, there are 232 TNLs, 39 CNLs, and 16 RNLs (Fig. 1e-f, Supplementary Table S2). It is known that the C-JID domain, a domain found in TNLs, is essential for the recognition of pathogen effectors (Martin et al., 2020). We identified 199 NLRs containing the C-JID domain in the Westar genome, which represents 43.2% of the whole TIR-contained proteins (Supplementary Table S3). Out of the 428 partial NLRs, 138 sequences were found to only carry the TIR domain.

In addition to the canonical NLRs, we identified the integrated domains of NLRs (NLR-ID) in the NLRome of Westar canola by querying the Pfam database using InterProScan (E-value < 1E-5). Integrated domains often act as decoys, resembling crucial host protein components targeted by pathogen effectors, and play a key role in initiating the defense response upon effector recognition. A total of 69 NLRs were found to carry integrated domains, and 49 different types of domains were identified (Supplementary Fig. S3 and Table S4). The galactose oxidase domain was found to be the most prevalent in 8 NLRs, with 3 tandemly duplicated NLRs carrying multiple copies of the domain (Supplementary Fig. S3 and Table S4). Other known integrated domains, such as the Heavy-Metal Associated domain (HMA), B3, Zinc-finger, and Kinase, were also present (Supplementary Fig. S3 and Table S4).

By providing a near-complete NLR repertory for *B. napus*, our study serves as a vital resource for the plant biotechnology community, fostering further research and application in crop species. For example, we are now able to identify inter- and -intra genomics duplication events of NLR-encoding genes between the subgenomes A and C of *B. napus* (Supplementary Fig. S4), something that was not possible with the wrongly annotated Westar canola genome. Moreover, compared to other NLRome studies, the Westar NLRome provides the first complete open reading frame, start and stop codons, which can give more information to researchers and breeders (Supplementary File S3). These findings are not only a significant advancement in understanding canola genetics but also offer practical applications for breeding programs and biotechnology aimed at improving disease resistance.

## ACKNOWLEDGEMENTS

The authors thank Yanick Asselin at Université Laval for the guidance during the RenSeq baits designing. This work was supported by the Saskatchewan Canola Development Commission (SaskCanola) and the Western Grains Research Foundation (WGRF) project Understanding the molecular basis of NLR-mediated clubroot resistance in *Brassica napus*. We are also very thankful to Fonds de recherche du Québec—Nature et technologies (FRQNT) for providing graduate funding to Jiaxu Wu.

## CONFLICT OF INTEREST

No conflict of interest is declared.

## AUTHOR CONTRIBUTIONS

E.P.L. managed the project; J.W. and S.M. performed the experiments and analyzed the data; J.W., S.M. and E.P.L. wrote the manuscript; J.W., S.M. and E.P.L. prepared figures.

## DATA AVAILABILITY STATEMENT

The raw data generated in this study is available in the Bioproject: PRJNA1137270, BioSample SAMN42564584, and SRA SRS22046306. The code used in this project can be found in Github link: https://github.com/Edelab. The rest of the data that supports the findings of this study are available in the supplementary material of this article.

## SUPPORTING INFORMATION

Additional supporting information may be found online in the Supporting Information section at the end of the article.

**Figure S1**. Expanded schematic representation of workflow followed in this study.

**Figure S2**. Frequency of the 345 NLR originally annotated in Westar genomes A and C by chromosomes.

**Figure S3**. Main NLR integrated domains identified in Westar genome.

**Figure S4**. Collinearity analysis of NLR genes between Westar A genome and C genome.

**Table S1**. Full list of NLRs and types of Westar genome.

**Table S2**. NLR numbers by chromosome and types.

**Table S3**. Full list of NLRs with C-JID domains generated by NLRtracker.

**Table S4**. Full list of NLRs with integrated domains.

**File S1**. Nucleotide sequences sent to Arbor Biosciences for baits design.

**File S2**. The predicted NLR loci annotated by NLR-Annotator.

**File S3**. Amino acid sequences of identified NLR proteins in this study.

## Notes

### Competing Interest Statement

The authors have declared no competing interest.

## REFERENCES

Alamery, S., Tirnaz, S., Bayer, P., Tollenaere, R., Chaloub, B., Edwards, D. and Batley, J. (2018) Genome-wide identification and comparative analysis of NBS-LRR resistance genes in Brassica napus. Crop and Pasture Science 69, 72–93.

Chen, X., Tong, C., Zhang, X., Song, A., Hu, M., Dong, W., Chen, F., Wang, Y., Tu, J., Liu, S., Tang, H. and Zhang, L. (2021) A high-quality Brassica napus genome reveals expansion of transposable elements, subgenome evolution and disease resistance. Plant Biotechnology Journal 19, 615–630.

Javed, M.A., Schwelm, A., Zamani-Noor, N., Salih, R., Silvestre Vañó, M., Wu, J., González García, M., Heick, T.M., Luo, C., Prakash, P. and Pérez-López, E. (2023) The clubroot pathogen Plasmodiophora brassicae: A profile update. Molecular Plant Pathology 24, 89–106.

Jupe, F., Witek, K., Verweij, W., Śliwka, J., Pritchard, L., Etherington, G.J., Maclean, D., Cock, P.J., Leggett, R.M., Bryan, G.J., Cardle, L., Hein, I. and Jones, J.D.G. (2013) Resistance gene enrichment sequencing (RenSeq) enables reannotation of the NB-LRR gene family from sequenced plant genomes and rapid mapping of resistance loci in segregating populations. The Plant Journal 76, 530–544.

Kourelis, J., Sakai, T., Adachi, H. and Kamoun, S. (2021) RefPlantNLR is a comprehensive collection of experimentally validated plant disease resistance proteins from the NLR family. PLOS Biology 19, e3001124.

Marchal, C., Michalopoulou Vassiliki A., Zou, Z., Cevik, V. and Sarris Panagiotis F. (2022) Show me your ID: NLR immune receptors with integrated domains in plants. Essays in Biochemistry 66, 527–539.

Martin, R., Qi, T., Zhang, H., Liu, F., King, M., Toth, C., Nogales, E. and Staskawicz, B.J. (2020) Structure of the activated ROQ1 resistosome directly recognizing the pathogen effector XopQ. Science 370, eabd9993.

Steuernagel, B., Witek, K., Krattinger, S.G., Ramirez-Gonzalez, R.H., Schoonbeek, H.-j., Yu, G., Baggs, E., Witek, A.I., Yadav, I., Krasileva, K.V., Jones, J.D.G., Uauy, C., Keller, B., Ridout, C.J. and Wulff, B.B.H. (2020) The NLR-Annotator Tool Enables Annotation of the Intracellular Immune Receptor Repertoire. Plant Physiology 183, 468–482.

Thomas, W.J.W., Amas, J.C., Dolatabadian, A., Huang, S., Zhang, F., Zandberg, J.D., Neik, T.X., Edwards, D. and Batley, J. (2024) Recent advances in the improvement of genetic resistance against disease in vegetable crops. Plant Physiology.

