## Supplementary figures and images for "Refinement of the *Brassica napus* NLRome using RenSeq"

### Fig. S1

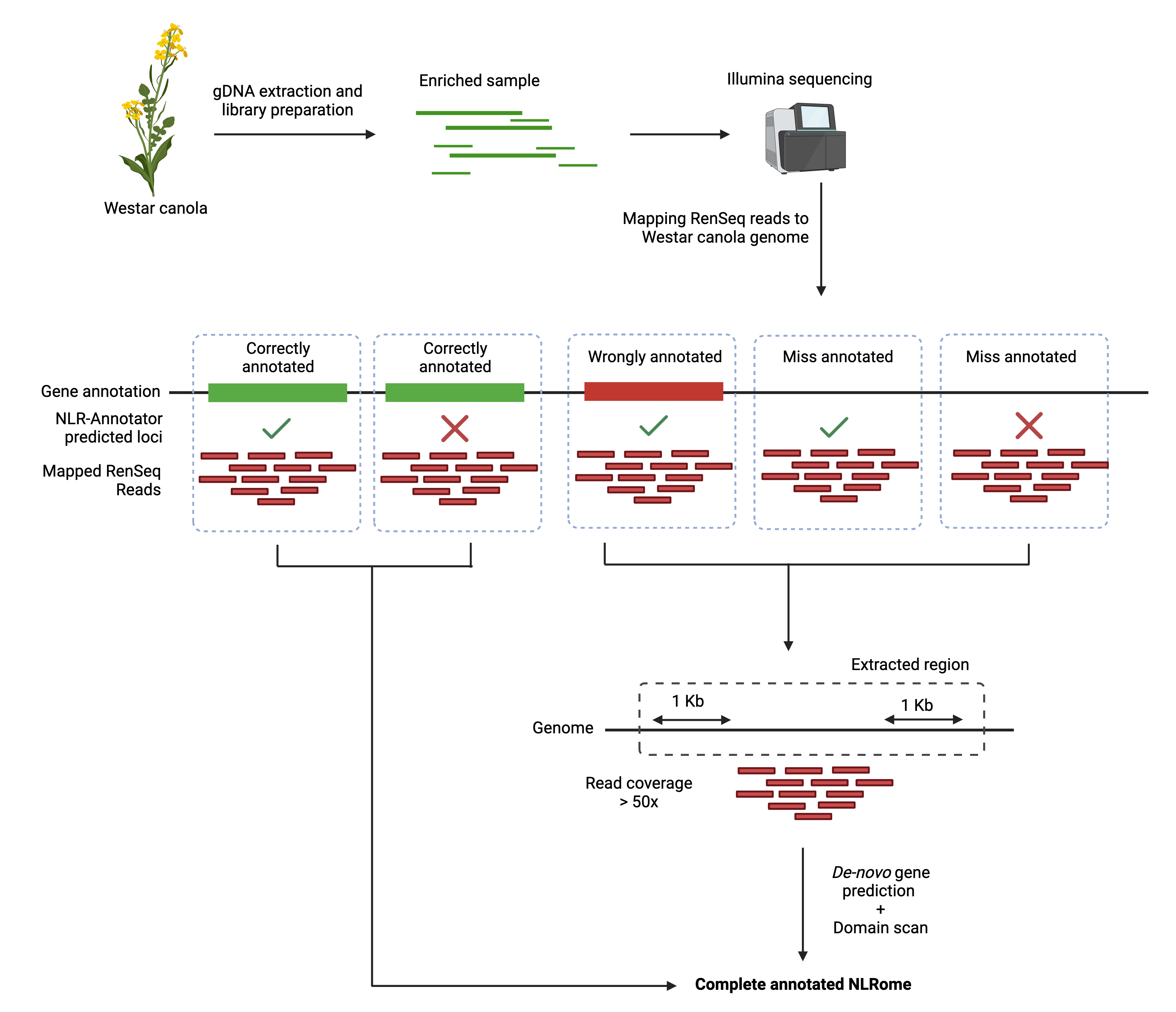

### Fig. S2

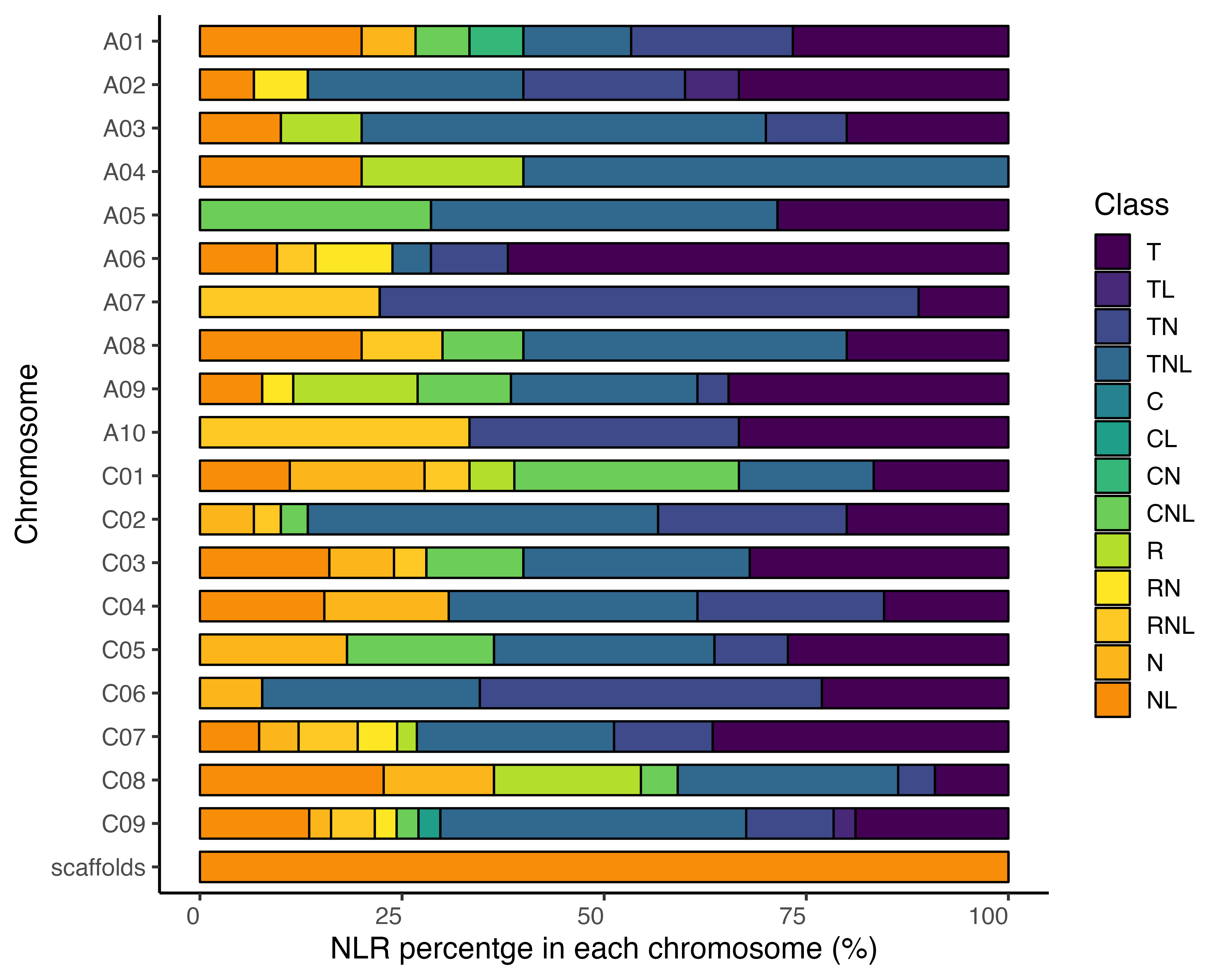

### Fig. S3

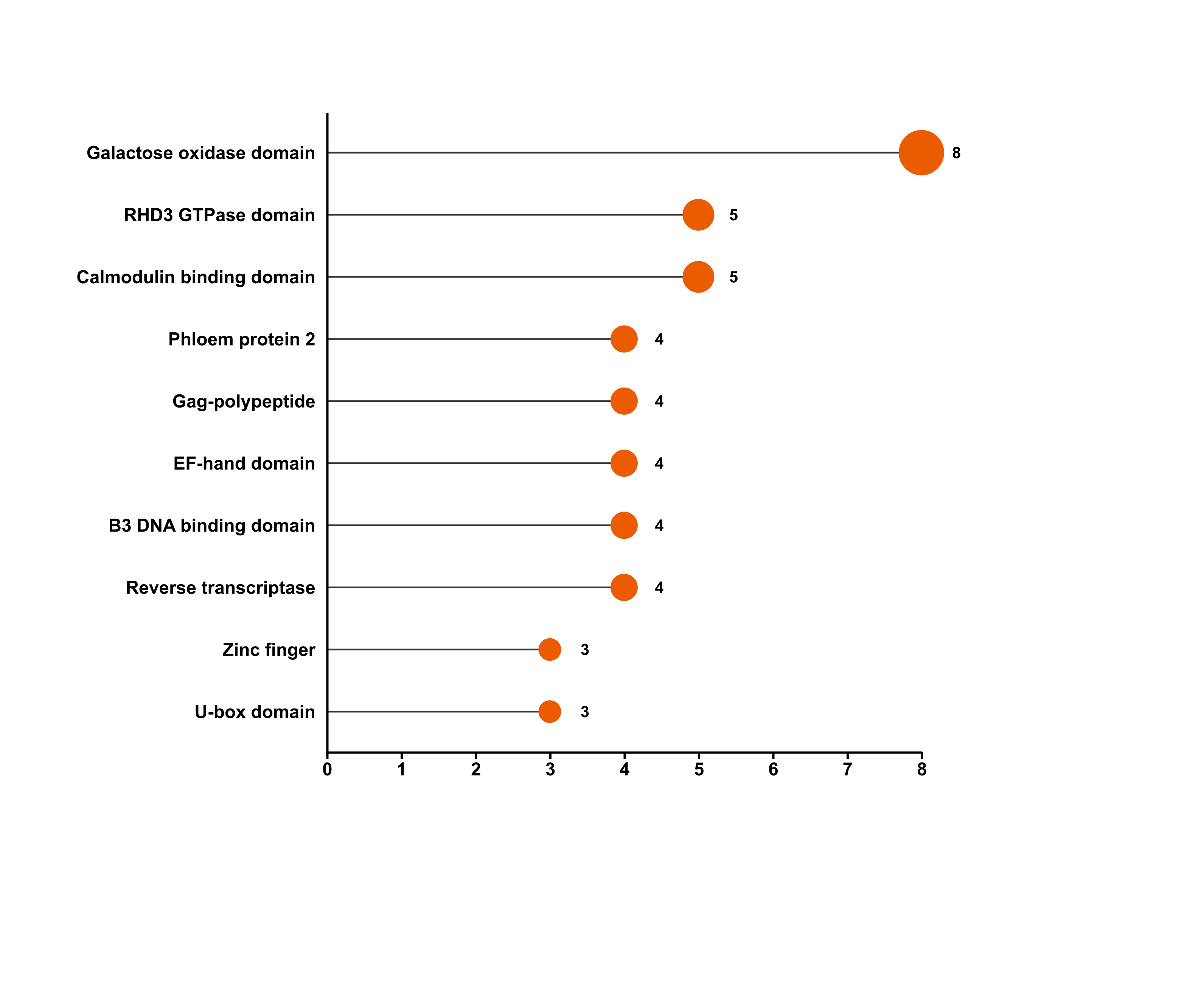

### Fig. S4

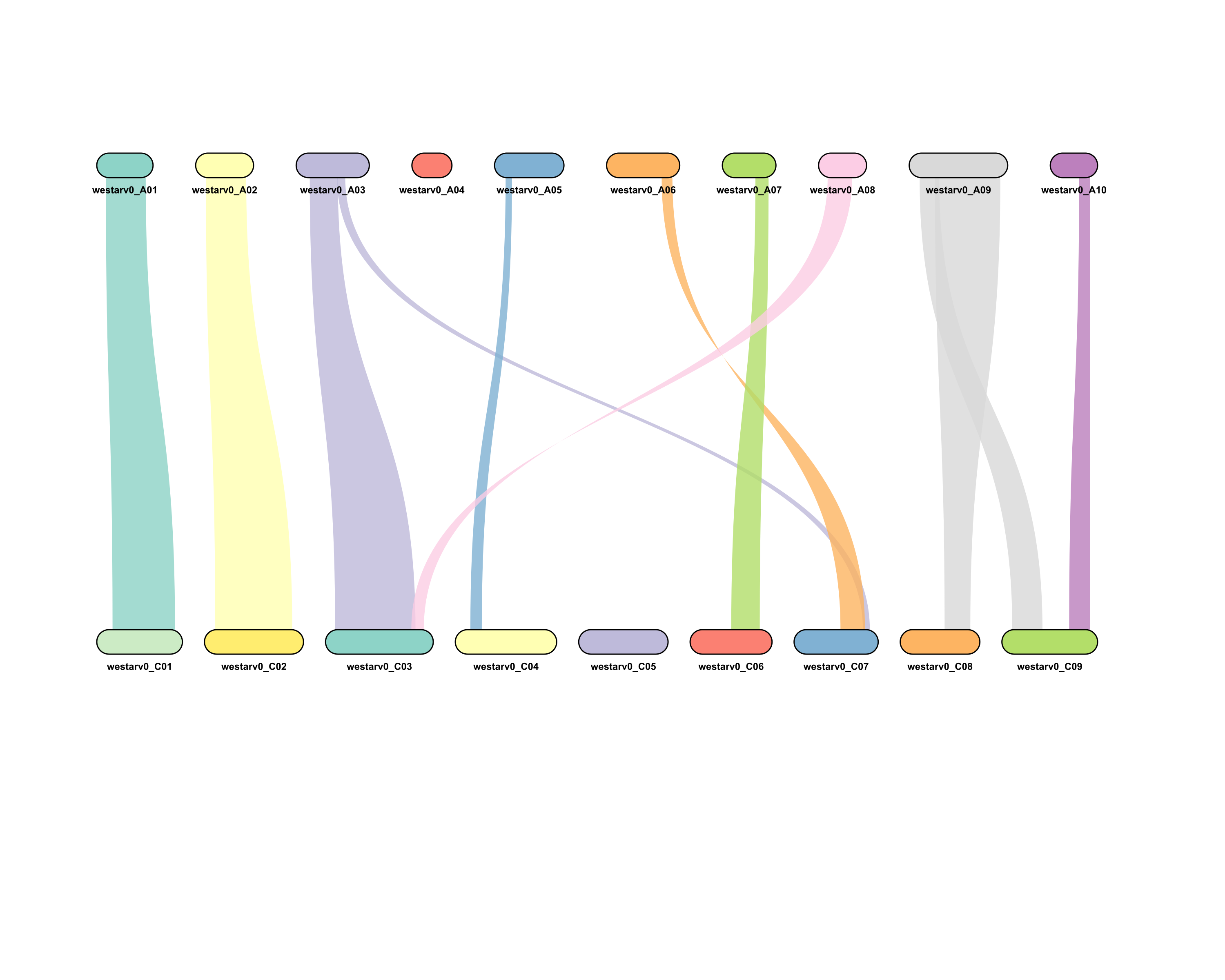
